# Climate change and intensive land use reduce soil animal biomass through dissimilar pathways

**DOI:** 10.1101/2020.01.26.920330

**Authors:** Rui Yin, Julia Siebert, Nico Eisenhauer, Martin Schädler

## Abstract

Global change drivers, such as climate and land use, may profoundly influence body size, density, and biomass of organisms. It is still poorly understood how these concurrent drivers interact in affecting ecological communities. We present results of an experimental field study assessing the interactive effects of climate change and land-use intensification on body size, density, and biomass of soil microarthropods. We found that both climate change and intensive land use decreased their total biomass. Strikingly, this reduction was realized via two dissimilar pathways: climate change reduced mean body size, while intensive land use decreased population size. These findings highlight that two of the most pervasive global change drivers operate via different pathways when decreasing soil animal biomass. These shifts in soil communities may threaten essential ecosystem functions like organic matter turnover and nutrient cycling in future ecosystems.

**Significance:** Many important ecosystem functions are determined by the biomass of soil animal, however, how their biomass may respond to climate change and land-use intensification still remains unknown. We conducted a large field study to investigate the potential interaction between these two pervasive global change drivers, and disentangle the pathways where they contribute to the changes in soil animal biomass. Our findings are exceptionally novel by showing detrimental, but largely independent, effects of climate change and land-use intensity on soil animal biomass, and that these independent effects can be explained by two dissimilar pathways: climate change reduced mean body size, while intensive land use decreased population size. Notably, consistent climate change effects under different land-use regimes suggest that (1) the identified pathways may apply to a wide range of environmental conditions, and (2) current extensive land-use regimes do not mitigate detrimental climate change effects on ecosystems.

## Introduction

Anthropogenic environmental changes are altering ecological communities and ecosystem functions (Chapin et al., 2000; Sala et al., 2000). As one of the most pervasive drivers, climate may change the functioning and evolutionary adaptations of communities (Briones et al., 2009; Hoffmann and Sgró, 2011). For instance, climate change can have substantial influences on population-level phenotypic trait expression of organisms, such as shifts in morphology, i.e. body size and shape (Gardner et al., 2011).

As warmer conditions increase individual metabolism (Scheffers et al., 2016) and development rates (Atkinson et al., 2003), many groups of organisms like plants, fish, ectotherms, birds, and mammals have already been reported to shrink their body size in response to warming (Sheridan and Bickford, 2011). These shifts in body size may result in a wide range of implications, e.g. biomass loss, including negative effects on the structure and dynamics of ecological networks (Woodward et al., 2005). Organisms with small body size in general and soil-living organisms in particular have not received much attention in this context, although a few studies on micro- and mesofauna in aquatic ecosystems revealed the same phenomena (Merckx et al., 2018). Although it is well known that the high biodiversity in soil drives a plethora of essential ecosystem functions (Bardgett and Van Der Putten, 2014; Hättenschwiler et al., 2005; Wall et al., 2015), it is still not clear how the body size, density, and biomass of soil organisms vary in response to climate change.

Besides climate change, soil systems further strongly suffer from land-use intensification, e.g. due to conversion of (semi-) natural areas into arable land or grasslands into croplands with intensified management practices (Foley et al., 2011). Such practices include tillage, mowing, livestock grazing, heavy machine employment, as well as herbicide, pesticide, and fertilizer application, all of which may profoundly imperil soil communities, and the functions and services they provide (Giller, 1997; McLaughlin and Mineau, 1995; Newbold et al., 2015; Tsiafouli et al., 2015). Accordingly, land-use change is considered as the major global threat for biodiversity (Sala et al., 2000), and this view also holds for soil ecosystems (Smith et al., 2016).

Soil biodiversity and food web structures have been reported to suffer from changes in land-use intensity (Mäder et al., 2002; Tsiafouli et al., 2015) and land-use conversion (e.g., from grasslands to croplands) (French et al., 2017). In this context, intensive land use is known to decrease the abundance and biodiversity of soil organisms (Bardgett and Van Der Putten, 2014; Flynn et al., 2009; Postma-Blaauw et al., 2010), consequently threatening the functioning of soils and the ecosystem services that they deliver, like soil fertility and nutrient dynamics (Beare et al., 1992; de Vries et al., 2013; Yin et al., 2019a), which may be fed back to primary production (Cardinale et al., 2004). Furthermore, intensive land use can have implications for trait diversity and the composition in above- and belowground arthropod communities (Birkhofer et al., 2017a). For example, frequent perturbations in intensive land use may select for soil microarthropods with particular life-history traits, such as *r*-strategists with high reproduction rates and small body size.

Taken together, both climate change and intensive land use may decrease the biomass of soil microarthropods by reducing their mean body size and population density. Such changes in soil communities would be alarming given that many important ecosystem functions are determined by the biomass of organisms in the soil (Höfer et al., 2001; Horwath, 1984; Petersen and Luxton, 1982). Moreover, climate change effects on soil communities and their functions can be dependent on environmental contexts, such as different land-use regimes (De Vries et al., 2012; Foley et al., 2005; Walter et al., 2013). That is, intensively-used land characterized by higher levels of disturbance and lower biodiversity may be more vulnerable to climate change (Isbell et al., 2017), while extensively-managed land with less disturbance and higher biodiversity potentially mitigate these detrimental effects of climate change (Oliver et al., 2016). Therefore, disentangling the pathways by which these main environmental change drivers contribute to changes in the biomass of soil organisms and identifying potential interactive effects is crucial to better understand how ecosystem functions and services may be affected and could be maintained in the future.

To address this critical knowledge gap, we tested potential interactive effects of climate and land use on body size, density, and biomass of soil microarthropods. This study was conducted at the Global Change Experimental Facility (GCEF; Fig. S1a) in Central Germany, where climatic conditions are manipulated following a future scenario for the years 2070-2100 with increased temperature (ambient vs. ∼0.6°C warming) and altered precipitation patterns (20% reduction in summer, and 10% addition in spring and autumn, respectively) across five different land-use regimes (two croplands and three grasslands differing in management intensity). We used data of multiple sampling campaigns and performed structural equation modeling to test different pathways how climate, land use, and the interaction of these two could affect the total biomass of soil microarthropods. We hypothesized that (1) climate change and intensive land use will decrease the body size and density of soil microarthropods, which then causes a reduction in total microarthropod biomass. Moreover, (2) we expected to find synergistic effects of these two environmental change drivers as detrimental climate change effects may be particularly strong in intensively-used land, by contrast, in extensively-used land these detrimental climate effects can be diminished.

## Materials and Methods

### Study site

The Global Change Experimental Facility (GCEF) is a large field research platform of the Helmholtz-Center for Environmental Research (UFZ), which is located in Bad Lauchstädt, Germany (51° 23’ 30N, 11° 52’ 49E, 116 m a.s.l.) and was established on a former arable field with the last cultivation in 2012. This arable field is characterized by a low mean annual precipitation of 498 mm and a mean temperature of 8.9°C. The soil is a Haplic Chernozem with neutral pH (5.8 – 7.5), high nutrient contents (i.e., total carbon and total nitrogen varied between 1.71 – 2.09% and 0.15 – 0.18%, respectively), and humus content of 2% reaching down to a depth > 40 cm. The soil is known for high water storage capacity (31.2%) and density (1.35 g/cm^3^), ensuring a relatively low sensitivity to drought stress (Altermann et al., 2005; WRB, 2007).

### Experimental set-up

The GCEF platform was designed to investigate the effects of future climatic conditions on ecosystem processes under different land-use regimes (Schädler et al., 2019). Each of the ten main-plots was divided into five sub-plots (16 m x 24 m), resulting in 50 sub-plots in total. The five sub-plots within each main-plot were randomly assigned to one of the five land-use regimes: (1) conventional farming, (2) organic farming, (3) intensively-used meadow, (4) extensively-used meadow, and (5) extensively-used pasture (grazed by sheep). Half of the main-plots are subjected to an ambient climate scenario, the other half to a future climate scenario (Fig. S1b, 1c).

Croplands and intensive meadows were established on the respective sub-plots in summer and autumn of 2013. The intensive meadow is a conventionally used mixture of forage grasses (20% *Lolium perenne*, 50% *Festulolium*, 20% *Dactylis glomerata*, and 10% *Poa pratensis*). Within the study period, winter wheat (2015) and winter barley (2016) were grown in these two croplands. In extensively-used meadow and pasture, we repeatedly sowed target plant seeds (legumes, grasses and non-leguminous dicots) during spring and autumn of 2014. For detailed description see Table S1.

**TABLE 1.** Results (F-values) from a repeated-measures ANOVA testing the effects of climate, land use, and their interactions on (a) body size, (b) density and (c) biomass of soil microarthropods, Acari, and Collembola. Significant effects are indicated in bold font, with † = *P* < 0.1, * = *P* < 0.05, ** = *P* < 0.01, *** = *P* < 0.001.

| Independent variable | *Df* | (a) Body size |  |  | (b) Density |  |  | (c) Biomass |  |  |
| --- | --- | --- | --- | --- | --- | --- | --- | --- | --- | --- |
|  |  | Microarthropods | Acari | Collembola | Microarthropods | Acari | Collembola | Microarthropods | Acari | Collembola |
| Climate (C) | 1,8 | **7.5\*** | 2.08 | **4.20<sup>†</sup>** | 1.96 | 1.81 | **3.65<sup>†</sup>** | **8.98\*** | **6.60\*** | **4.54<sup>†</sup>** |
| Land use (L) | 4,32 | 0.59 | 1.47 | 0.90 | **6.68\*\*\*** | **10.1\*\*\*** | 1.22 | **4.15\*\*** | **4.36\*** | 1.18 |
| C × L | 4,32 | 1.47 | 2.34 | 0.58 | 0.74 | 1.11 | 0.2 | 1.16 | 1.39 | 0.14 |

The climate treatment is based on a consensus scenario for Central Germany in the period from 2070 to 2100, which was derived from 12 climate simulations based on four different emission scenarios using three established regional climate models: COSMO-CLM (Rockel et al., 2008), REMO (Jacob and Podzun, 1997), and RCAO (Döscher et al., 2002). The consensus scenario predicts an increase of mean temperature across all seasons by 1 to 2°C. For precipitation, mean values of the 12 projections resulted in an experimental treatment consisting of a ∼9% increase in spring (March – May) and autumn (September – November) and a ∼21% decrease in summer (June – August).

All main-plots are equipped with steel framework elements (5.50 m height) to account for possible side effects of the construction itself. Main-plots that are subject to future climate are further equipped with mobile shelters, side panels, and rain sensors to allow for alterations in precipitation amounts. Shelters were automatically closed from sundown to sunrise to increase night temperature (Beier et al., 2004). The night closing during these periods increased the mean daily air temperature at 5 cm-height by 0.55°C, as well as the mean daily soil temperature in 1 cm- and 15 cm-depth by 0.62°C and 0.50°C, respectively. By using an irrigation system, we added rain water to achieve ∼110% of ambient rainfall to the main-plots with future climate in spring and autumn. Additionally, the rain sensors associated with the irrigation system were used to regulate precipitation on the future climate main-plots to ∼80% of ambient rainfall in summer. As a result, precipitation was increased by 9.2% to 13.6% in spring and autumn and decreased by 19.7% to 21.0% in summer in both years, respectively. Climate manipulation started in spring 2014. During our experiment, the roofs were active from 15^th^ February – 11^th^ December in 2015 and from 22^nd^ March to 29^th^ November in 2016.

### Assessment of soil microarthropods

Soil samples were collected three times (autumn 2015, spring 2016, and autumn 2016) during a 1.5-year study period. At each sampling point, three soil cores (Ø 6 cm, 5 cm depth) were taken per sub-plot to extract microarthropods (Collembola and Acari) using a Macfadyen high-gradient extraction method (Macfadyen, 1961). Using a digital microscope (VHX-600, Keyence Corp., Osaka, Japan), Acari were identified to order level, i.e., Oribatida, Mesostigmata and Prostigmata; and Collembola were identified to family level, i.e., Isotomidea, Entomobryidae, Katiannidae, Sminthurididae, Hypogastruridae and Onychiuridae. For all taxa we counted the number of individuals, and measured the body size (length, μm) of each individual using the measurement function of the VHX microscope.

### Statistical analysis

The biomass (M, μg) of microarthropod groups (Order level: Collembola, Oribatida, Mesostigmata and Prostigmata) was calculated according to a specific formula: Log M = a + b × log L with L as the body size (length) of microarthropods (μm), with Collembola: a = - 1.8479; b = 2.3002; Mesostigmata: a = 2.064; b = 2.857; Oribatida: a = 2.117; b = 2.711; Prostigmata: a=2.124; b=2.808 (Ganihar, 1997; Mercer et al., 2001). For each sub-plot, the mean body size, population density, and total biomass of microarthropods, Acari and its orders (Oribatida, Mesostigmata, and Prostigmata), Collembola and its families (Isotomidea, Entomobryidae, Katiannidae, Sminthurididae, Hypogastruridae and Onychiuridae) were represented by Mean ± SD and analysed.

A repeated-measures split-plot ANOVA was conducted using a generalized linear mixed model (GLMM) with Type III sum of squares (procedure MIXED, SAS University Edition v9.4) to analyze these response variables as affected by the experimental treatments: climate (2 levels) was analyzed at the main-plot factor and land use (5 levels) and its interaction with climate were tested at the sub-plot factor. While these effects represented the between-subject model, the within-subject model accounts for the covariance structure of the residuals due to repeated measures (repeated measures with 3 sampling events). Post-hoc Tukey’s HSD tests were carried out to reveal significant differences among means.

Furthermore, we ran structural equation models using the *piecewiseSEM* package (Lefcheck, 2016) to disentangle the pathways by which climate and land use affect the total biomass of soil microarthropods (Fig. 2a). By using *piecewiseSEM*, we were able to account for the hierarchical study design, to test for interaction effects, and to include random effects (Lefcheck, 2016). More precisely, we employed structural equation modeling to test if climate, land use, and their interaction affected soil microarthropod biomass (main response variable) *via* reductions in body size and/or population density and if the two global change drivers differ in their pathways (Fig. 2a). The models were created using linear mixed-effects models within the *nlme* package (Pinheiro J, Bates D, DebRoy S, 2017) and tested the interactive effects of climate and land use on the body size and population density of microarthropods as well as the effects of body size and population density on the total biomass of microarthropods. A random intercept with main-plots nested within date was incorporated in the models. The list of models included the following:

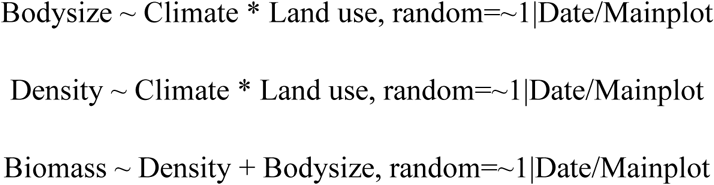

We reduced the number of variables using the Akaike Information Criterion (AIC), which is implemented in the *piecewiseSEM* package. A minimum change of 2 AIC units was used as a prerequisite for each simplification step. Further, we reported the standardized coefficient for each path of both models (i.e. the full conceptual model and the reduced final model) (Table S2, Fig. 2a, b). The overall fit of the models was evaluated by using Shipley’s test of d-separation obtained through Fisher’s C statistic (Table S3, Fig. 2b). The statistical analyses for the structural equation modeling were performed using the R statistical software (R Core Team, 2017).

## Results

### Climate change reduces the body size of soil microarthropods

Climate change significantly reduced the body size of microarthropods by ∼10% (Fig. 1a; Table 1a), which was mainly driven by the reduced body size of Collembola but not Acari (Fig. 1a, 1b; Table 1a). Specifically, the body size of dominant Collembola families i.e., Isotomidae, Entomobryidae and Sminthurididae significantly reduced under future climate scenario (Fig. S2a-c; Table S4a), by contrast, the body size of Acari orders did not profoundly respond to climate change, only Oribatida and Mesostigmata tended to decrease their body size (Fig. S2d-e; Table S4a).

**Figure 1.**
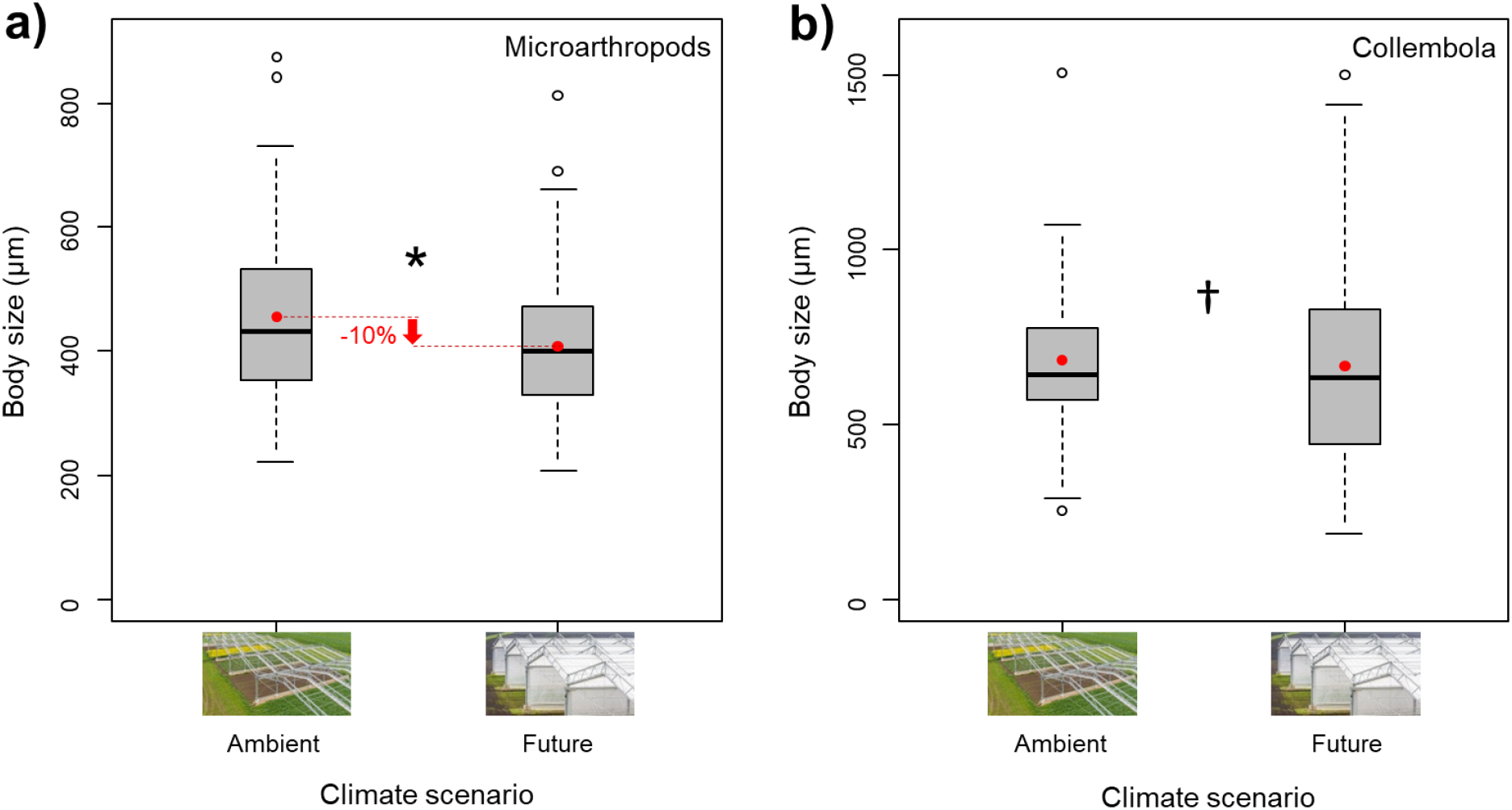
Effects of climate on body size of microarthropods (a) and Collembola (b). * and ^†^ denote significant (*P* < 0.05) and marginal (*P* < 0.10) differences between climate scenarios, respectively. Boxplots show the median (horizontal line), the mean (red dot), first and third quartile (rectangle), 1.5 × interquartile range (whiskers), and outliers (isolated white dots).

### Intensive land use reduces the density of soil microarthropods

Land-use intensification dramatically decreased the density of microarthropods by ∼47% from the extensively-used meadow to conventional farming (Fig. 2a; Table 1b), which was driven by the decreased density of Acari (Fig. 2b; Table 1b). All observed Acari orders, i.e., Oribatida, Mesostigmata and Prostigmata decreased their population density in response to land-use intensification from meadows to croplands (Fig. S3a-c; Table S4b). However, climate effects on microarthropods were negligible, specifically, future climate only marginally decreased the density of Collembola (Fig. 2c; Table 1b), which was caused by a significant decrease in the density of Isotomidae (Fig. S3d; Table S4b).

**Figure 2.**
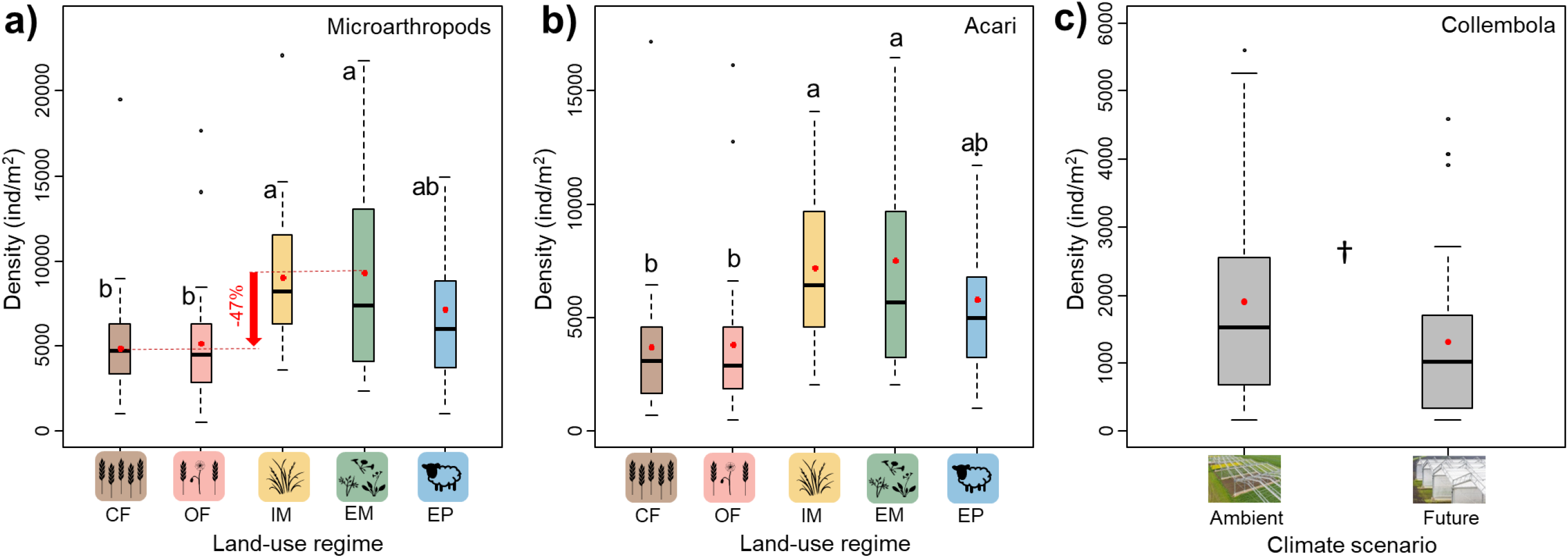
Effects of land use on density of microarthropods (a) and Acari (b), and effects of climate on density of Collembola (c). Different lowercase letters denote significant (*P* < 0.05) differences among land-use types based on Post hoc Tukey’s HSD tests. ^†^ represents marginal (*P* < 0.01) differences between climate scenarios. Boxplots show the median (horizontal line), the mean (red dot), first and third quartile (rectangle), 1.5 × interquartile range (whiskers), and outliers (isolated black dots). CF = conventional farming; OF = organic farming; IM = intensively-used meadow; EM = extensively-used meadow; EP = extensively-used pasture.

### Climate change and intensive land use reduce the biomass of soil microarthropods

The total biomass of microarthropods was significantly affected by both global change factors (Table 1). Specifically, climate change significantly reduced the biomass of microarthropods by ∼17% (Fig. 3a; Table 1c). The same decreasing pattern was found for the biomass of Acari (Fig. 3b; Table 1c), specifically, the total biomass of dominant Acari orders, i.e., Oribatida and Mesostigmata significantly decreased under future climate scenario (Fig. S4a-b; Table S4c); whereas the total biomass of Collembola was only marginally decreased by future climate (Table 1c), which was driven by a significant decrease in total biomass of Isotomidae (Fig. S4c; Table S4c). Additionally, the total biomass of microarthopods sharply decreased by ∼37% from the extensively-used meadow to conventional farming (Fig. 3d; Table 1c), which was driven by the decreased biomass of Acari (Fig. 3e; Table 1c), specifically, the total biomass of dominant Acari orders, i.e., Oribatida and Mesotigmata significantly decreased in response to land-use intensification from grasslands to croplands (Fig. S4d-e; Table S4c)..

**Figure 3.**
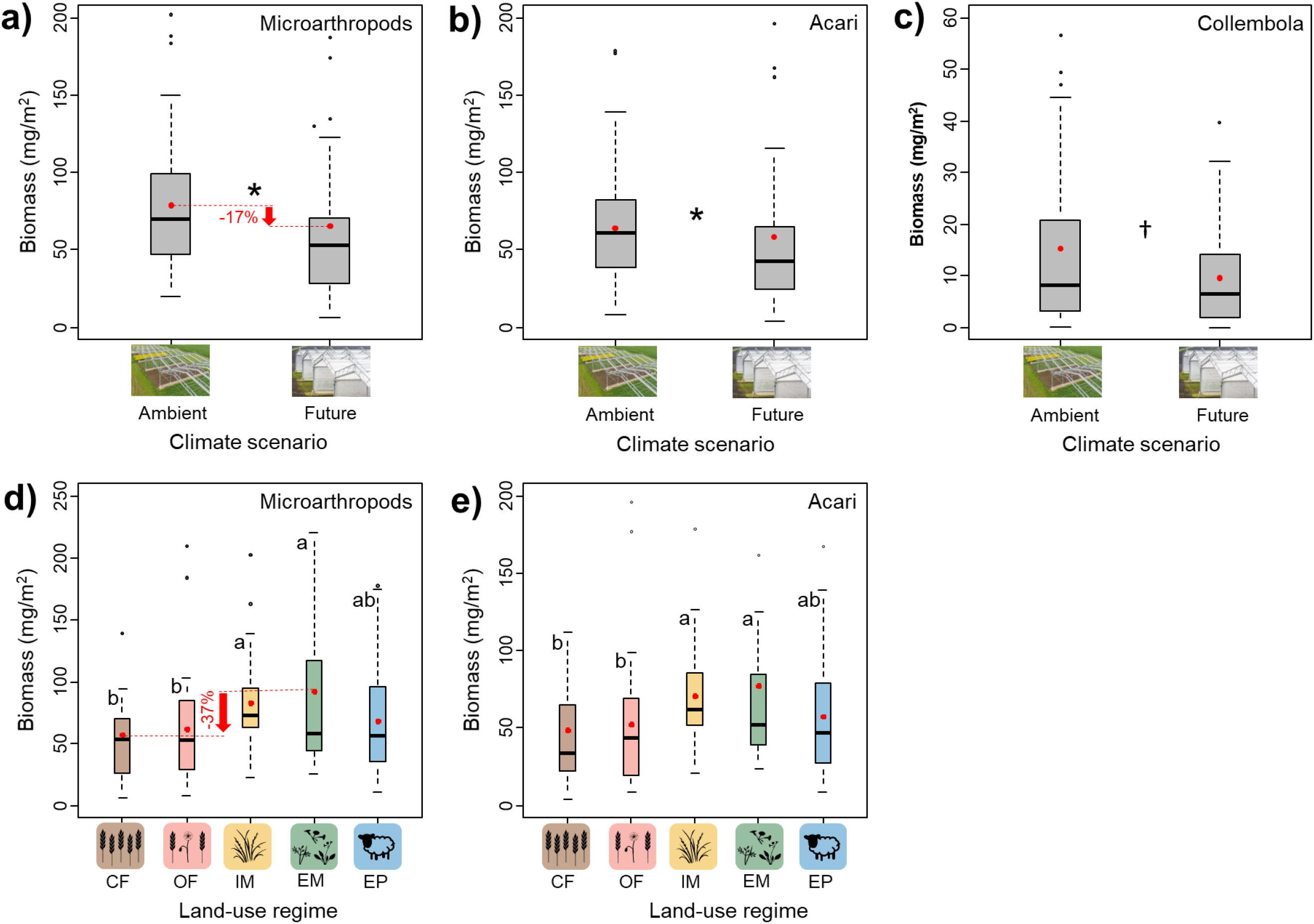
Effects of climate on biomass of microarthropods (a), Acari (b) and Collembola (c), and effects of land use on biomass of microarthropods (d) and Acari (e). * and ^†^ denote significant (*P* < 0.05) and marginal (*P* < 0.01) differences between climate, respectively. Different lowercase letters denote significant differences (*P* < 0.05) among land-use types based on by Post hoc Tukey’s HSD tests. Boxplots show the median (horizontal line), the mean (red dot), first and third quartile (rectangle), 1.5 × interquartile range (whiskers), and outliers (isolated black dots). CF = conventional farming; OF = organic farming; IM = intensively-used meadow; EM = extensively-used meadow; EP = extensively-used pasture.

**Figure 4.**
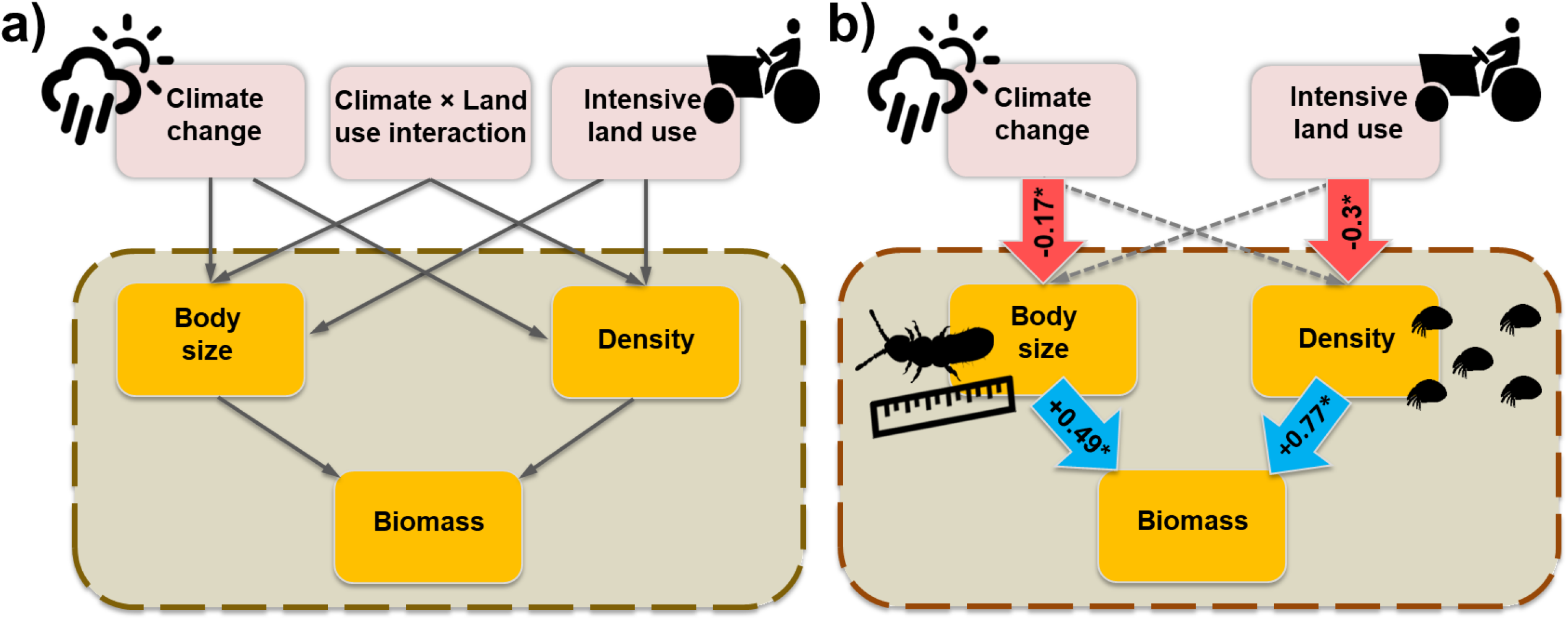
Structural equation model (SEM) showing the pathways through which climate change and land use influence soil microarthropod biomass. **(a)** A-priori model of hypothesized indirect climate and land -use effects and their interaction on the biomass of soil microarthropods across all sampling points. **(b)** Final selected model (AIC 39.02) with blue (positive) and red (negative) arrows indicating significant effects (*P* < 0.05). Dashes arrows indicate non-significant (*P* > 0.05) effects, but still remained in the final model. Fisher’s C = 3.016; *P* = 0.56; *d.f.* = 4. Numbers along the arrows are standardized path coefficients.

### No interactive effects of climate and land use on soil microarthropods

Contrary to our expectation, there were no significant interactive effects of climate and land use on body size, density, or biomass of soil microarthropods; neither on Acari nor Collembola (Table 1).

### Pathways of biomass decrease in soil microarthropods

SEM results revealed that climate change and intensive land use reduced the biomass of soil microarthropods *via* two different pathways. While biomass loss caused by future climate was mediated by reduced body size, biomass loss caused by intensive land use was mediated by decreased density (Fig. 2b, Table S2).

## Discussion

The main findings of the present study are that (1) climate change reduced the mean body size of soil microarthropods; (2) land-use intensification decreased their population density; and (3) these two pathways independently contributed to an overall reduction of total soil microarthropod biomass. As climate change and land-use intensification operated *via* dissimilar pathways, our hypothesis of synergistic environmental change effects was not supported, indicating that the underlying pathways of climate change effects are consistent across land-use regimes and vice versa.

### Climate change and land-use intensification reduce total microarthropod biomass

In our study, climate change caused a significant reduction of total microarthropod biomass in soil. Our results are in line with those of Vestergård *et al*. (2015), who demonstrated that the biomass of microarthropods was dramatically reduced by drought, especially when combined with warming. Accordingly, climate warming was shown to exacerbate the drying of soil and thereby the negative drought effects on soil microarthropods (Thakur et al., 2018).

For the soil system, we could show with our study that the future climate treatment simulated by increased air and soil temperatures (+0.6°C) and altered precipitation (−20% in summer and +10% in spring/autumn) consistently reduced the body size and total biomass of soil microarthropods across different land-use types and management. This adds to the existing body of literature reporting similar effects for other groups of organisms (Daufresne et al., 2009; Gardner et al., 2011; Sheridan and Bickford, 2011; Yom-Tov, 2001) and environmental contexts, thus underlining its universal validity.

Reduction in body size is supposed to be a universal response of animals to climate change, which is supported by the ecological rules dealing with temperature–size relationships, i.e., Bergmann’s rule (Bergmann, 1848), James’ rule (James, 1970), and Temperature–size rule (Atkinson, 1994), stating that warmer conditions would lead to organisms with smaller body size (Gardner et al., 2011). This phenotypical plasticity is one of the most taxonomically widespread patterns in biology (Forster et al., 2012). It’s worth noting that the effects on the reduced body size with increased temperatures are more profound in Collembola communities than Acari communities in our study. The projected future climate significantly decreased the body size of total Collembola and their most dominant families, i.e., Isotomidae, Entomobryidae and Sminthurididae; but the effects on the body size of total Acari were less, only showing the tendency to decrease in Oribatida and Mesostigamata. This finding might be supported by some other studies, which have shown that Collembola were more susceptible to warming than Acari, and Acari generally would take longer time to respond to climate change (Makkonen et al., 2011; Vestergård et al., 2015; Yin et al., 2019b).

Given the tight connection between ecological properties (e.g., longevity, fecundity and mortality rates, as well as competitive interactions) and body size (Chown and Gaston, 2010; Savage et al., 2004; Thakur et al., 2017), shifts in body size may have profound implications for ecosystem functioning, food web stability, and mass acquisition. For example, shrinking body size has often been shown to result in reductions in total biomass (e.g., decreased yield in fisheries and crops) (Sheridan and Bickford, 2011; Woodward et al., 2005).

Land-use intensity also reduced the total biomass of microarthropods, but this negative effect was mainly due to lower densities in croplands than in grasslands. This is in accordance with other studies reporting negative effects of intensive land use (Baker, 1998; Birkhofer et al., 2017b) on the density of soil fauna, whereas the conversion from croplands to grasslands was shown to have positive effects (Zaitsev et al., 2006). Lower densities of soil fauna in croplands might be due to multiple detrimental effects, i.e., vegetation composition and mono-cropping, mechanical disturbance of the upper soil horizon, application of agrochemicals, and exposure to desiccation. In contrast, grasslands with less extensive management practices, and more diverse plant communities, which provide an environment with more habitats and more accessible food sources, can maintain higher densities of soil organisms (Alvarez et al., 2001; Nyawira Muchane, 2012; Scherber et al., 2010). Accordingly, we found significantly higher levels of microarthropod biomass in grasslands than in croplands, which is supported by previous studies (e.g., de Groot et al., 2016).

### Extensive land use has limited potential to mitigate the consequences of climate change

It is often suggested that extensive land use may effectively mitigate climate change effects due to higher (above- and below-ground) diversity and lower anthropogenic disturbance (Isbell et al., 2017; Oliver et al., 2016). Accordingly, we hypothesized the detrimental effects of climate change on soil microarthropods could be particularly strong in intensive land use, whereas they could be compensated by extensive land use. In contrast to this hypothesis, we did not observe any significant interaction effects of climate and land use on body size, density, and total biomass of soil microarthopods. These results showed that the effects of climate change on soil microarthropods were consistent across different land-use regimes, suggesting that detrimental climate change effects will not be exacerbated by intensive land use, nor mitigated by extensive land use. This finding calls for novel management strategies to alleviate the consequences of climate change. Our study provides the first mechanistic insights into the underlying pathways of changes in soil communities that may inform such novel approaches.

### Climate change and land-use intensification decrease soil microarthropod biomass via independent pathways

Structural equation modelling revealed that climate change and intensive land use reduced the total biomass of soil microarthropods via two dissimilar and independent pathways: while climate change reduced the mean body size, intensive land use decreased overall densities. This is the first empirical evidence for such contrasting pathways underlying different environmental change factors in a full-factorial experiment. Consistent climate change and land-use effects under different land-use regimes and climate contexts, respectively, suggest that the identified pathways may apply to a wide range of environmental conditions. However, we do not know yet if the outcomes are due to pure assembly mechanisms or if evolution is also involved.

Additionally, body size-mediated effects of climate change on soil microarthropod communities may have profound implications for total community composition and ecosystem processes driven by soil organisms, such as by decreasing litter decomposition rates (Taylor et al., 2010). We therefore encourage future studies to investigate how microarthropod biomass shifts affect soil ecological processes (like litter decomposition and nutrient dynamics) and food web relations in the context of climate change. Moreover, future studies should investigate if decreasing mean body size of soil microarthopods in response to climate change is due to species turnover towards smaller *r*-selected species, shrinking body size within species, or both. In contrast, the reduction in microarthopod densities in soil due to land-use intensification was not accompanied by reductions in mean body size. Future studies should therefore explore which other traits of soil microarthropods are influenced by climate change and land-use intensity and how they link to ecosystem functioning.

## Conclusion

The results confirm our hypothesis of detrimental effects of climate change and intensive land use on the biomass of soil microarthropods. In contrast to our expectation, however, these two environmental change drivers reduced soil microarthropod biomass through dissimilar pathways, which may explain why more extensive land use did not attenuate detrimental climate change effects. Additionally, the reduced total biomass of soil microarthropods may have a wide range of ecological implications, as their biomass is known to be linked to several ecosystem functions and services (e.g., litter decomposition and nutrient cycling) and is therefore likely to trigger indirect effects of climate and land use on soil ecosystem functioning (Woodward et al., 2005). We encourage future studies to explore the links between these observed soil community changes and soil ecological processes as well as to explore novel management options that increase the population sizes, and thus total biomass, of soil microarthropods to mitigate climate change effects on the functioning of soils.

## Acknowledgments

Rui Yin as the first author appreciates the funding by the Chinese Scholarship Councils (CSC) (File No.201406910015). All authors appreciate the Helmholtz Association, Federal Ministry of Education and Research, the State Ministry of Science and Economy of Saxony-Anhalt and the State Ministry for Higher Education, Research and the Arts Saxony to fund the Global Change Experimental Facility (GCEF) project. This project also received support from the European Research Council (ERC) under the European Union’s Horizon 2020 research and innovation program (grant agreement No. 677232 to N.E.). Further support came from the German Centre for Integrative Biodiversity Research (iDiv) Halle-Jena-Leipzig, funded by the DFG (FZT 118). We also appreciate the staff of the Bad Lauchstädt Experimental Research Station (especially Ines Merbach and Konrad Kirsch) for their hard work in maintaining the plots and infrastructures of the GCEF, and Dr. Stefan Klotz, Prof. Dr. Francois Buscot and Dr. Thomas Reitz for their roles in setting up the GCEF. In addition, particularly thank Prof. Dr. Paul Kardol for his polishing of the manuscript and his valuable comments.

## Author Contributions

R.Y. and M.S. performed field work, R.Y., N.E. and M.S. conceived and planned the paper, R.Y. and J.S. performed the statistical analyses. All authors contributed to the interpretation of results and wrote the paper.

## SUPPLEMENTARY MATERIALS

**Table S1.** Descriptions of five land-use regimes.

| **Land-use regime**<br>(Abbreviation) | **Descriptions** |
| --- | --- |
| **Conventional farming**<br>(CF) | A typical regional crop rotation consisting of winter rape, winter wheat and winter barley with the application of mineral fertilizers and pesticides. |
| **Organic farming**<br>(OF) | A crop rotation aiming to maintain soil fertility, minimize measures of pest and weed control, and provide an environmentally friendly management of agro-ecosystems with mechanical weed control, organic fertilization, non-stained seeds and restricted use of pesticides. |
| **Intensively-used meadow**<br>(IM) | Conventional used mixture of forage grasses with moderate fertilization and frequent mowing (3-4 times per year). |
| **Extensively-used meadow**<br>(EM) | A wide range of native common grasses, herbs and legumes (totally consisting 50 plant species, for each species seeds were sampled from different local populations to reflect to local gene pool and to introduce genetic variability) with moderate mowing (2-3 times per year) and no fertilization. |
| **Extensively-used pasture**<br>(EP) | Plant species composition and land management as the same as extensively used meadows (see above), but with sheep grazing (2-3 grazing periods per year with a group of 20 sheep grazing for 24 hours). |

**Table S2.** The standardized coefficient for each path in the reduced final model.

| Response variable | Explanatory variable | Std. Estimates | P-value |
| --- | --- | --- | --- |
| Body size | Climate | **-0.170** | **0.026** |
|  | Land use | 0.112 | 0.060 |
| Density | Climate | -0.078 | 0.301 |
|  | Land use | **-0.281** | **< 0.001** |
| Biomass | Body size | **0.490** | **< 0.001** |
|  | Density | **0.766** | **< 0.001** |

**Table S3.** The overall fit of the models. Marginal R²: model variation explained by fixed effects. Conditional R²: model variation explained by both fixed and random effects.

| Response variable | R<sup>2</sup> (marginal) | R<sup>2</sup> (conditional) |
| --- | --- | --- |
| Body size | 0.04 | 0.60 |
| Density | 0.08 | 0.29 |
| Biomass | 0.59 | 0.62 |

**Table S4.** Results (F-values) from a repeated-measures ANOVA testing the effects of climate, land use, and their interactions on **(a)** body size, **(b)** density, and **(c)** biomass of Acari order (i.e., Oribatida, Mesostigmata, Prostigmata) and Collembola family (i.e., Isotomidae, Entomobryidae, Katiannidae, Sminthurididae, Hypogastruridae, Onychiuridae). Significant effects are indicated in bold font, with † = *P* < 0.1, * = *P* < 0.05, ** = *P* < 0.01, *** = *P* < 0.001.

| Independent variable | (a) Body size |  |  | (b) Density |  |  | (c) Biomass |  |  |
| --- | --- | --- | --- | --- | --- | --- | --- | --- | --- |
| | Climate (1,8) | Land use (4,32) | C × L (4,32) | Climate (1,8) | Land use (4,32) | C × L (4,32) | Climate (1,8) | Land use (4,32) | C × L (4,32) |
| **<u>Acari</u>** |  |  |  |  |  |  |  |  |  |
| Oribatida | **3.54<sup>†</sup>** | 0.69 | 2.14<sup>†</sup> | 1.74 | **10.1\*\*\*** | 1.11 | **3.75\*** | **4.49\*\*** | 1.19 |
| Mesostigmata | **2.38<sup>†</sup>** | 2.19<sup>†</sup> | 1.23 | 0.67 | **6.99\*\*\*** | 1.29 | **7.06\*** | **2.24<sup>†</sup>** | 0.75 |
| Prostigmata | 0.46 | 0.56 | 0.85 | 3.18 | **8.09\*\*\*** | 0.54 | 1.67 | 0.97 | 0.43 |
| **<u>Collembola</u>** |  |  |  |  |  |  |  |  |  |
| Isotomidae | **3.63\*** | 1.14 | 1.57 | **7.69\*** | 1.63 | 0.94 | **5.73\*** | 0.27 | 0.48 |
| Entomobryidae | **4.83\*** | 1.66 | 2.89 | 1.8 | 1.1 | 0.19 | 0.14 | 0.68 | 1.02 |
| Sminthurididae | **3.7\*** | 0.5 | 1.61 | 2 | 0.68 | 0.38 | 0.23 | 0.47 | 0.25 |
| Katiannidae | 0.42 | 0.55 | 2.48 | 0.51 | 0.54 | 0.64 | 0.1 | 0.5 | 0.38 |
| Hypogastruridae | 0.52 | 0.32 | 1.06 | 1.53 | 1.41 | 2.27 | 0.16 | 0.18 | 0.42 |
| Onychiuridae | 0.08 | 0.76 | 0.32 | 0.04 | 0.24 | 0.48 | 0.05 | 0.62 | 0.4 |

**Figure S1.**
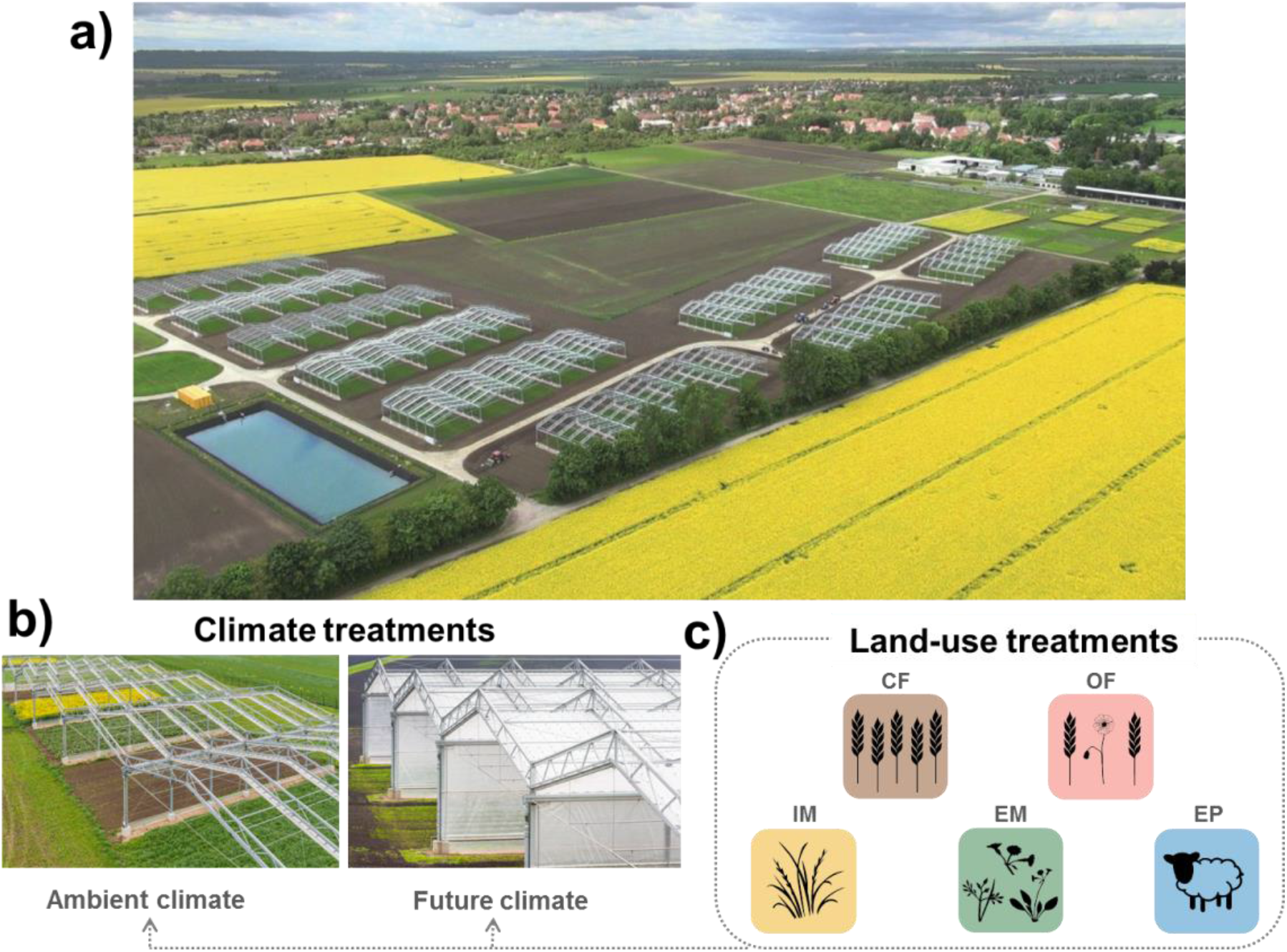
Global Change Experimental Facility (GCEF). (a) Aerial image of the experimental set-up of the GCEF in Bad Lauchstädt, Germany. Picture: Tricklabor/Service Drone, copyrights: UFZ. (b) Climate treatments as mainplot factor with two levels (ambient climate vs. future climate). Picture: Andrè Künzelmann, copyrights: UFZ. (c) Land-use treatments as sub-plot factor with five land-use regimes, abbreviations: CF = conventional farming; OF = organic farming; IM = intensive-used meadow; EM = extensive-used meadow; EP = extensive-used pasture.

**Figure S2.**
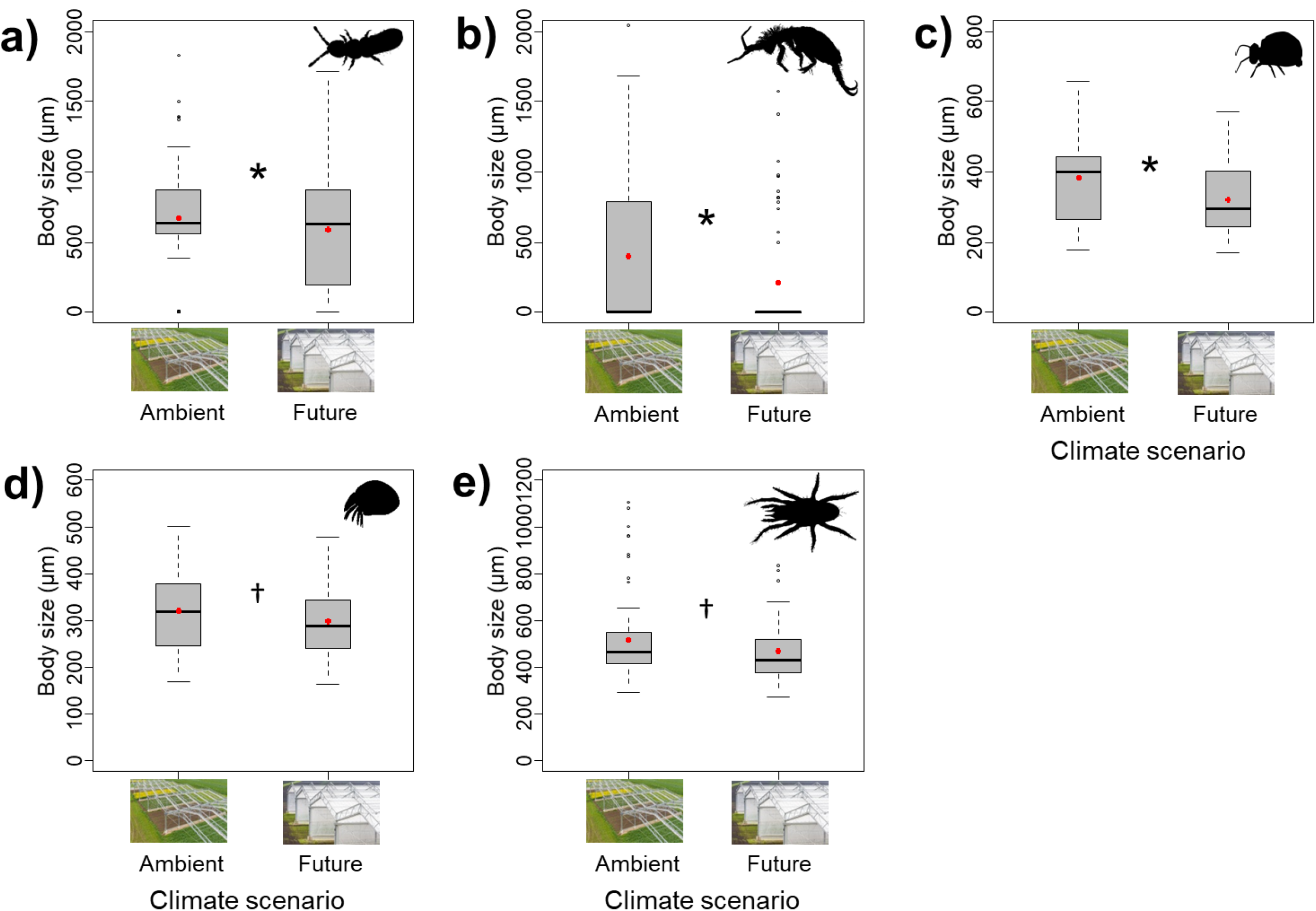
Effects of climate on body size of Isotomidae (a), Entomobryidae (b), Sminthurididae (c), Oribtida (d) and Mesostigmata (e). * and ^†^ denote significant (*P* < 0.05) and marginal (*P* < 0.10) differences between climate scenarios, respectively. Boxplots show the median (horizontal line), the mean (red dot), first and third quartile (rectangle), 1.5 × interquartile range (whiskers), and outliers (isolated white dots).

**Figure S3.**
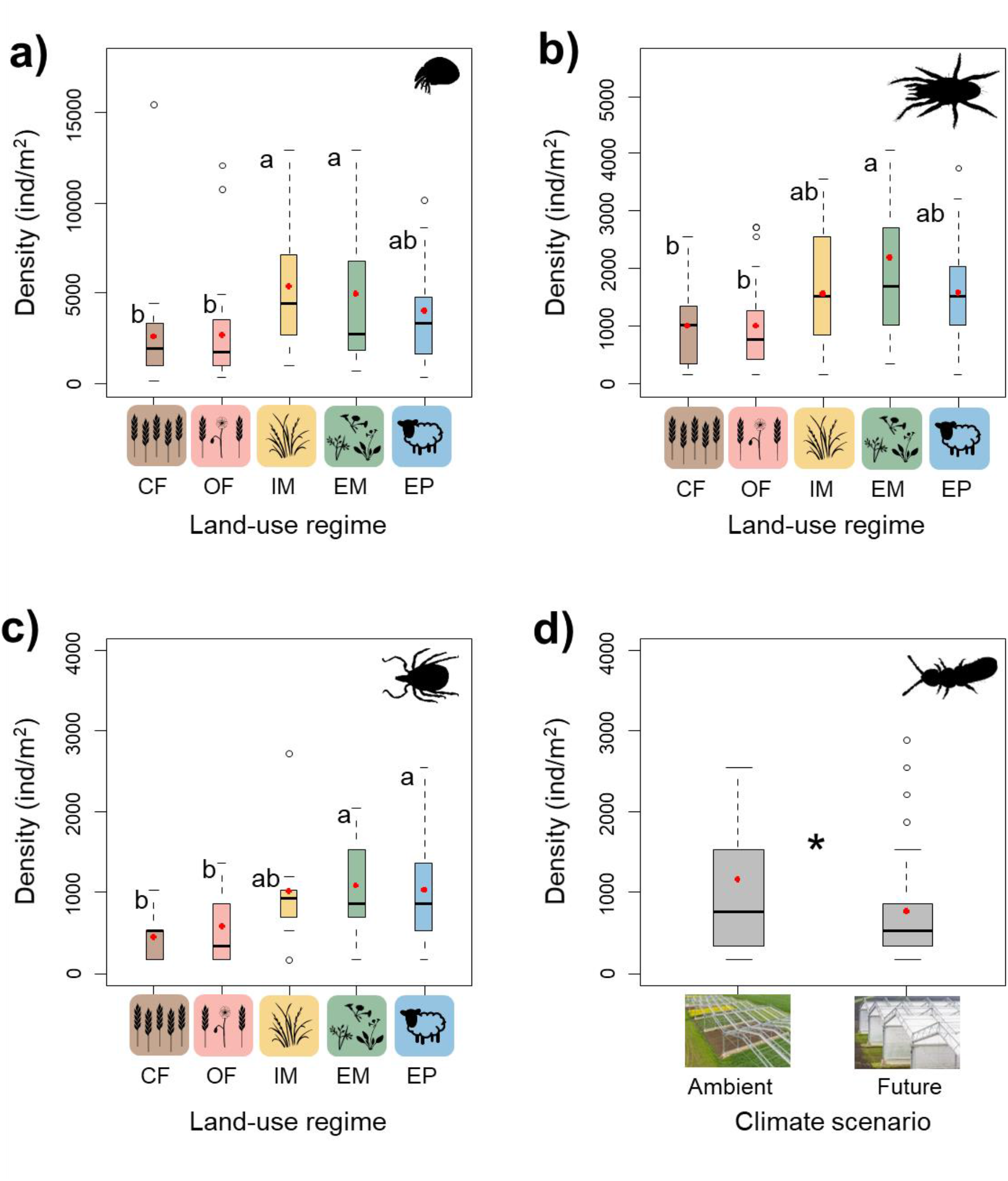
Effects of land use on density of Oribatida (a), Mesostigmata (b) and Prostigmata (c), and effects of climate on density of Isotomidae (d). * denote significant (*P* < 0.05) differences between climate, respectively. Different lowercase letters denote significant differences (*P* < 0.05) among land-use types based on by Post hoc Tukey’s HSD tests. Boxplots show the median (horizontal line), the mean (red dot), first and third quartile (rectangle), 1.5 × interquartile range (whiskers), and outliers (isolated black dots). CF = conventional farming; OF = organic farming; IM = intensively-used meadow; EM = extensively-used meadow; EP = extensively-used pasture.

**Figure S4.**
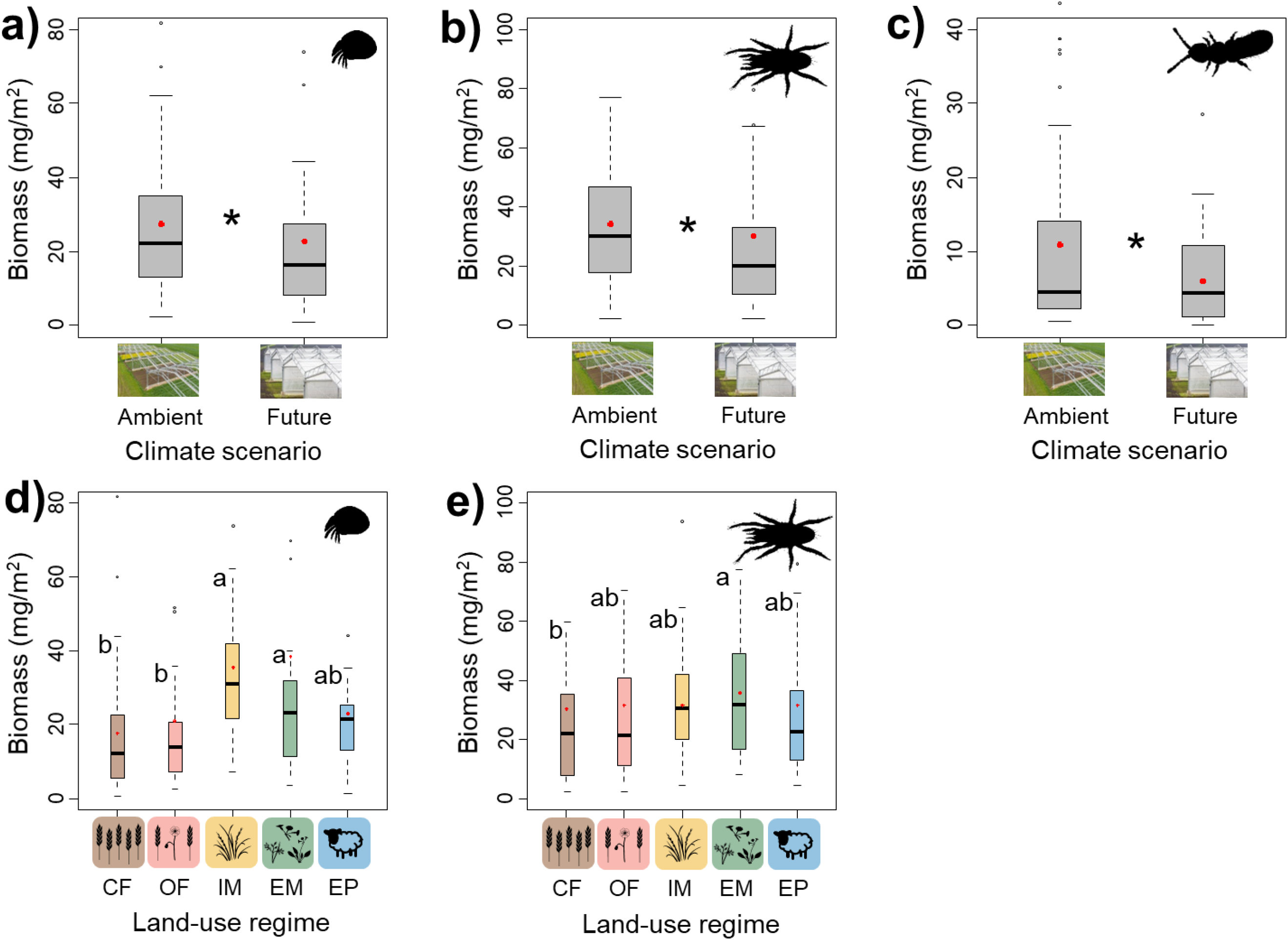
Effects of climate on biomass of Oribatida (a), Mesostigmata (b) and Isotomidae (c), and effects of land use on density of Oribatida (d) and Mesostigmata (e). * denote significant (*P* < 0.05) differences between climate, respectively. Different lowercase letters denote significant differences (*P* < 0.05) among land-use types based on by Post hoc Tukey’s HSD tests. Boxplots show the median (horizontal line), the mean (red dot), first and third quartile (rectangle), 1.5 × interquartile range (whiskers), and outliers (isolated black dots). CF = conventional farming; OF = organic farming; IM = intensively-used meadow; EM = extensively-used meadow; EP = extensively-used pasture.

